# Transplanted hearts assimilate the recipient’s biological age

**DOI:** 10.64898/2026.09.15.751836

**Authors:** Jesse R. Poganik, Tomohisa Matsunaga, Alexander Tyshkovskiy, Ake Lu, Amin Haghani, Hao Zhou, Friederike Martin, Steve Horvath, Michael M. Givertz, Stefan G. Tullius, Vadim N. Gladyshev

## Abstract

**Background:** Despite the growing demand for transplantation, a shortage of donor organs persists, limiting access to this lifesaving procedure. To meet the increasing demand, organs are frequently transplanted across an age discrepancy between donor and recipients.

**Methods:** We delineated biological age dynamics in heterochronic heart transplants performed in both mice and patients. We carried out multiomic profiling and applied advanced biomarkers of aging to understand the interactions between the transplanted organ and the host.

**Results:** We show that the biological age of experimental heart transplants rapidly assimilates the age of recipients. Interestingly, this effect was limited to the grafted tissue without reciprocal effects on systemic biological age of the recipient. These effects were confirmed at the level of DNA methylation and gene expression. Extending our findings clinically using both omics and functional analyses, we show that the biological age of heterochronically transplanted hearts is strongly associated with the age of the recipient, rather than the donor, confirming the clinical relevance of our experimental findings.

**Conclusions:** These data identify the systemic environment as a driving force on tissue biological age. The rejuvenation of older organs in young recipients suggests novel approaches to organ allocation that may expand the pool of transplantable organs.

## Introduction

Heart transplantation (hTx) is being performed across the world with increasing frequency, driven in large part by the growth of an aging population.^1^ In the United States alone, the number of recipients 65 years of age and older increased by 127% between 2010–2021.^2^ Unfortunately, the unprecedented demand for this lifesaving procedure has been accompanied by a persistent inability to meet the need for organs with the current donor pool. As a consequence, hundreds of patients are dying each year while waiting for a heart.^3^ This tragic reality has led to ongoing interest in methods to expand the pool of transplantable hearts.

Hearts for transplantation are selected based on an expansive list of criteria, including chronological age. Although no official upper limit exists, in practice donors younger than 45 years of age are recommended,^4^ and few transplant programs accept hearts from donors older than 50, although this limit has been and continues to be raised. As such, heterochronic transplantations, i.e. those in which donor and recipient age differ, are increasingly being performed to meet the demand.^5^

Organ transplantation offers a unique platform to investigate aging biology as it uniquely enables delineating effects of grafting a tissue of a particular age into an environment of differing age, an interaction of relevance observed as early as 1966.^6^ Compelling results from classical heterochronic parabiosis models have recently shown that biological age of an organism may be modulated by an age-disparate environment, with exposure to a youthful physiological environment resulting in rejuvenation and lifespan extension.^7^

Conversely, older environments may increase biological age.^8^ As transplantation involves exposure of an organ to an age-mismatched systemic environment, comparable biological age modulations may occur. For instance, previous work has shown that transplantation of old hearts in mice promotes the accumulation of senescent cells in tissues of the recipient, suggesting the exchange of aging features between donors and recipients.^9,10^

Here, we characterize the effects of donor and recipient age-mismatched heart transplantation using advanced biomarkers based on DNA methylation (DNAm) and transcriptomic profiling. Using a mouse model of heterotopic heart transplantation, we show that cardiac transplants rapidly assimilate to the age of recipients, with younger hearts grafted into older animals exhibiting accelerated aging, and younger recipient age initiating rejuvenation of grafted old hearts. Biological age modulation was limited to grafted tissue, and native tissues were not affected by the transplant. Extending these data to human transplant recipients, we documented similar effects of age assimilation, most notably with older hearts in younger recipients undergoing marked rejuvenation. Analysis of functional outcomes from hundreds of recipients one-year post-transplant confirmed the clinical relevance of these findings. These data provide a “real world” example of tissue rejuvenation with important implications for organ transplantation.

## Results

### Heterochronic heart transplantation in mice strongly affects the age of syngeneic cardiac transplants

Utilizing an established mouse model of syngeneic heterotopic cardiac transplant in which the recipient mouse retains its native heart, and a second heart is anastomosed to the carotid artery and jugular vein (**Figure 1A**), we performed a series of transplantations using donors and recipients of varying ages: 3 months (young, “Y”), 1 year (middle-aged, “M”), and 1.5–1.67 years (old, “O”). Heterochronic transplants were performed, in which the age of the donor and recipient diverged, and isochronic transplants, with donors and recipients of the same age (see Methods for details). To focus solely on the effects of aging, syngeneic transplants were performed between inbred C57Bl/6 to exclude the impact of alloimmune responses. Animals that underwent sham surgery were also analyzed. Four to six months post-transplantation, mice were euthanized and tissues were collected. We profiled DNAm using the HorvathMammalMethylChip320 array, which measures methylation levels at approximately 320,000 CpG sites relevant to mice, and applied DNAm biomarkers of aging. We analyzed transplanted hearts to delineate effects of the systemic environment on the graft tissue; in parallel, we analyzed native hearts, livers, and blood samples to probe systemic feedback of the graft.

**Figure 1.**
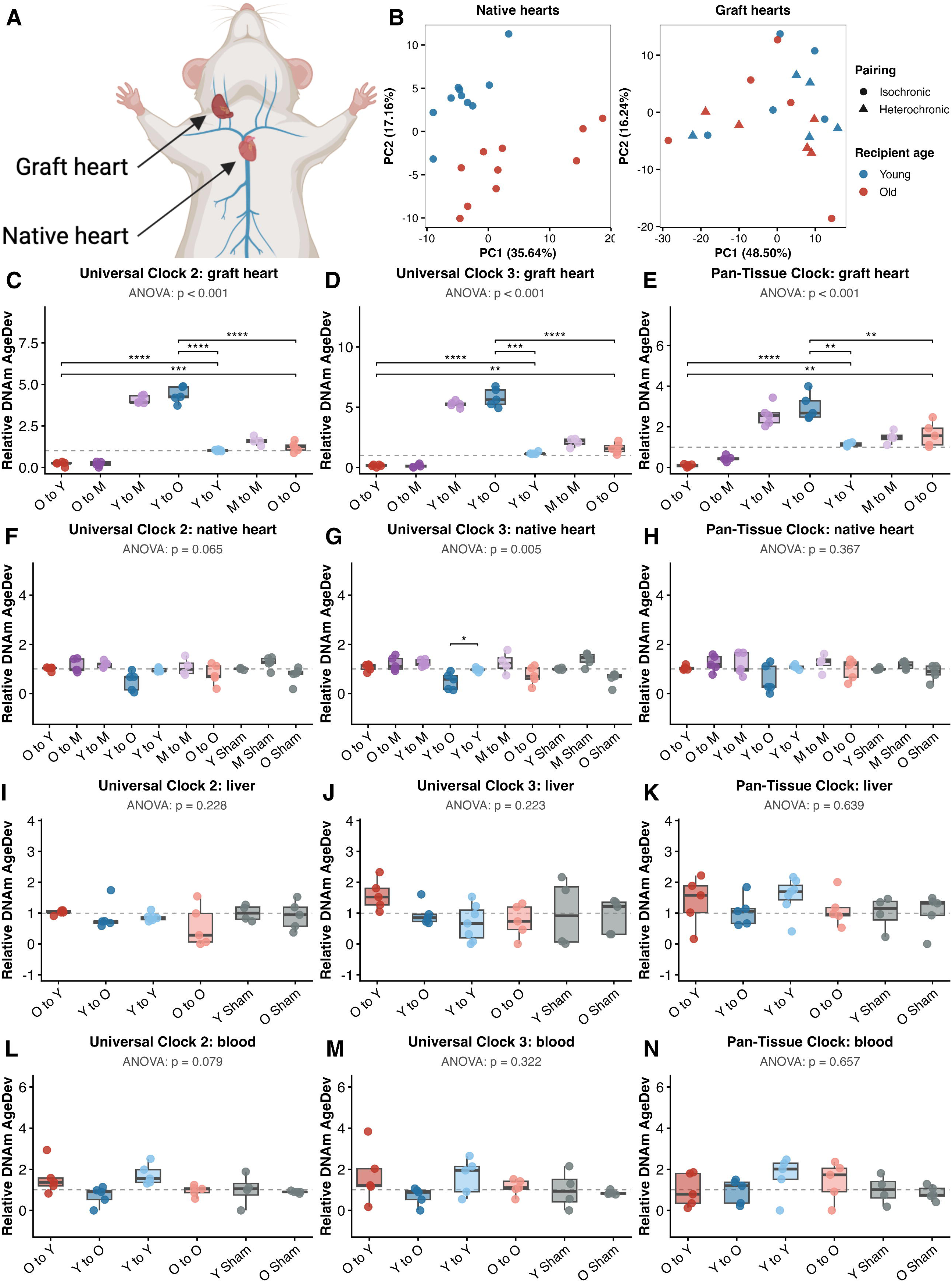
DNAm biomarkers of aging reveal biological age assimilation of hearts transplanted heterochronically with minimal impact on systemic biological age. (A) Schematic of heterotopic mouse transplant model. (B) Principal component analysis of DNAm data, computed separately for native hearts and graft hearts. (C–E) Relative DNAm AgeDev for grafted hearts. (F–H) Relative DNAm AgeDev for native hearts. (I–K) Relative DNAm AgeDev for livers. (L–N) Relative DNAm AgeDev for blood. Data are shown for Universal Clock 2 (C,F,I,L), Universal Clock 3 (D,G,J,M) and the Horvath pan-tissue clock (E,H,K,N). P values were calculated with ANOVA and unpaired t-tests. Sample sizes: M to M graft and native hearts, n = 4; Y sham native heart, n = 4; for all other groups, n=5. Y, young; M, middle-aged; O, old.

To assess whether age-related methylation differences were detectable at the genome-wide level, we performed principal component analysis (PCA) separately on native and transplanted hearts. Although the first principal component did not show a consistent association with recipient age, donor age, sex, or transplant group, chronological age was clearly captured along the second principal component in native hearts. Transplanted hearts, by contrast, showed no comparable separation (**Figure 1B**). This finding suggests that while normal chronological aging produces a broad, genome-wide methylation signature detectable by unsupervised analysis, the effect of the heterochronic environment on graft methylation may reflect a narrower set of aging-associated changes. We therefore investigated whether a biomarker-based analysis could better define age-related changes in the graft hearts.

We chose to analyze three DNAm aging biomarkers in this dataset, first applying two universal pan-mammalian, pan-tissue clocks, which were trained on DNAm data from 185 mammalian species: (i) “Universal Clock 2”, which predicts age relative to the maximum lifespan of a given species; and (ii) “Universal Clock 3”, which predicts age relative to the age of sexual maturity of a given species.^11^ We also analyzed a pan-tissue clock specific to mice.^12^ These biomarkers were selected because they are applicable to all tissues tested; training of the underlying models was robust; and they have been independently validated in relevant settings, e.g. heterochronic parabiosis.^7,8^ Moreover, we found strong, significant correlations between predicted age and chronological age in animals subjected to sham surgery, where we expect minimal perturbations of DNAm age (**Figure S1**). Age deviation (AgeDev), the difference between the predicted age and chronological age, was used as the basis of statistical comparison across chronological age groups.^13^

We first analyzed transplanted hearts. Based on all clocks applied, DNAm age of transplanted hearts was significantly affected by the age of the recipient: younger hearts grafted into older recipients were predicted older than the donor age, while older hearts implanted into younger recipients were predicted younger than the donor age (**Figure 1C–E; Table S1**). In contrast, the AgeDev of transplanted hearts in isochronic recipients was not affected as strongly, though we noted generally increased AgeDev in these groups (compared to native hearts), suggesting that transplantation itself somewhat increases DNAm age. This effect may be due to the physiological stress of the surgical procedure.^8^

We next assessed feedback of the grafted heart onto the native tissues of recipient mice. Analysis of the native hearts revealed no consistent significant effects, though we noted a slight reduction of AgeDev in native hearts with young transplants in old recipients (**Figure 1F–H; Table S1**). DNAm age of recipient blood and livers was similarly unaffected by heterochronic hTx (**Figure 1I–N; Table S1**). Thus, donor/recipient age-disparate heart transplantation significantly affects the age of the heart transplant while not showing strong systemic feedback on the recipient.

### DNAm patterns at the CpG and gene levels indicate accelerated aging in young heterochronic grafts and rejuvenation in old heterochronic grafts

To confirm our results using DNAm biomarkers of aging, we examined differential methylation levels of individual CpG sites. We identified 475 CpG sites that were significantly differentially methylated (adjusted p value < 0.05 and Δβ > 5%) in young heterochronic grafts compared to young isochronic controls, with 324 sites featuring increased methylation and 151 with decreased methylation (**Figure 2A** **and Table S2**). Conversely, old heterochronic grafts were found to have fewer differentially methylated sites overall compared to isochronic controls (51 with increased methylation, 62 with decreased methylation; **Figure 2B**). Next, we assessed changes that could be ascribed to aging by comparing old and young isochronic heart transplants and identified 161 and 184 sites with significantly increased and decreased methylation, respectively (**Figure 2C**). Interestingly, native hearts of old compared to young sham controls were found to have the most differential methylation (909 and 1,641 featuring increased and decreased methylation; **Figure 2D**), capturing a chronological aging signal in the absence of any transplant-related manipulation.

**Figure 2.**
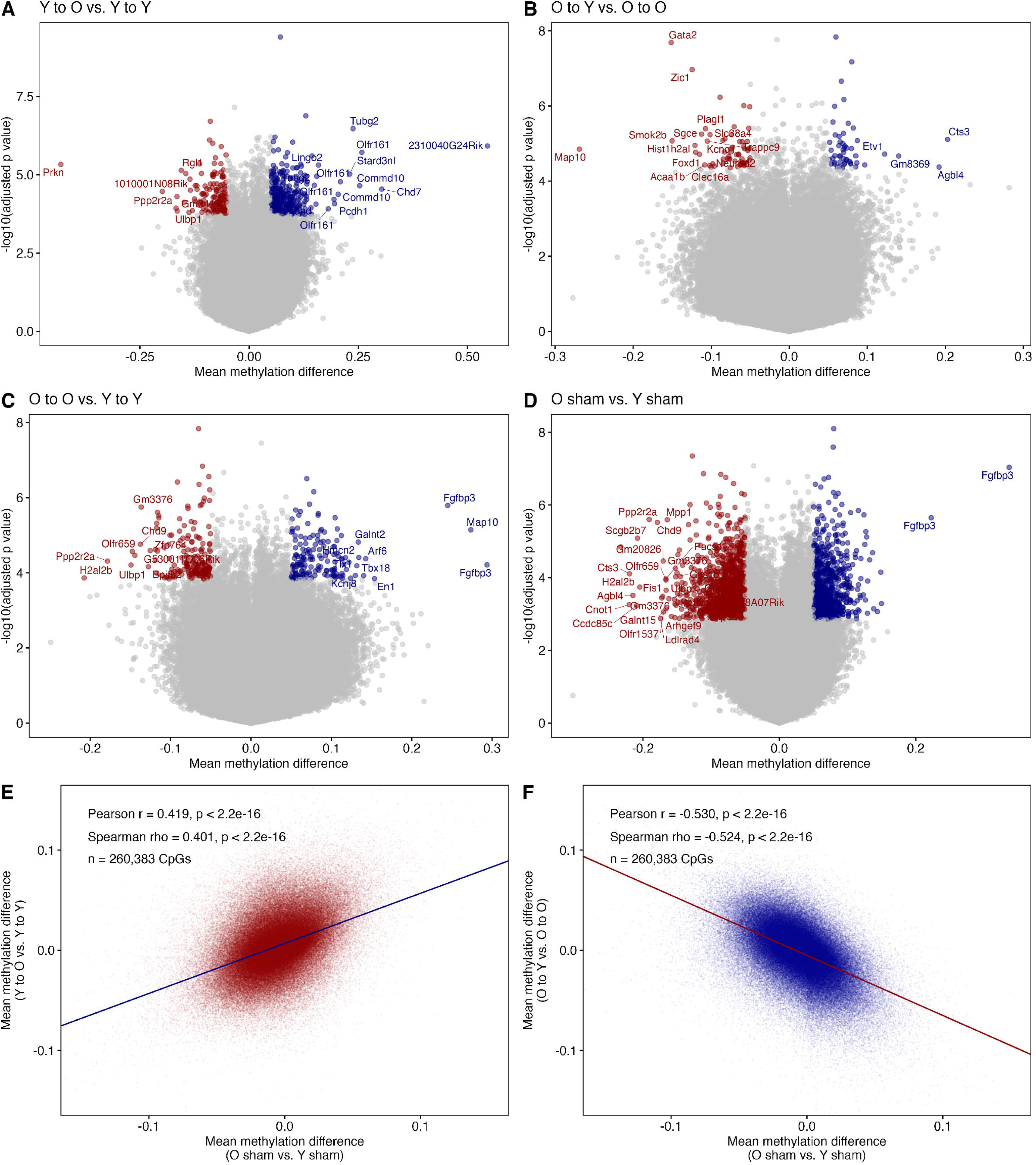
DNAm changes at the CpG site level support biological age assimilation in heterochronic heart grafts. (A–D) Volcano plots showing differential methylation analysis for young heterochronic compared to young isochronic heart transplants (A), old heterochronic compared to old isochronic heart transplants (B), Old isochronic vs. young isochronic grafts (C), and native hearts of old sham animals vs. young sham animals (D). Dots represent identified CpG sites and are labeled with their associated gene name. Colored dots represent significantly differentially methylated sites (adjusted p value < 0.05 and Δβ > 5%). (E–F) Correlation between methylation changes in heterochronic heart grafts and aging-associated methylation changes, assessed genome-wide across all measured CpG sites without restriction to a pre-defined subset, for young (E) and old heterochronic transplants (F), each compared against an independent aging reference (native hearts of old versus young sham animals). Pearson correlation and associated p value is shown. For E–F, dots represent individual CpG sites labeled with their associated gene name. Sample sizes: Y sham native heart, n = 4; n = 5 for all other groups. Y, young; O, old.

To assess whether these transplant-induced methylation changes reflected the direction of aging-associated methylation change, we correlated methylation changes across all measured CpG sites genome-wide, without restricting our analysis to any pre-defined subset between heterochronic grafts and an independent comparison of aging (old versus young sham animals). Consistent with our biomarker analysis, methylation changes in young hearts grafted into old environments were correlated positively with aging-associated changes (**Figure 2E**). Most strikingly, this correlation was inverted for old hearts grafted into young environments (**Figure 2F**). Thus, aging-associated methylation patterns are reversed in old hearts grafted into young environments, supporting the notion that these hearts were rejuvenated upon heterochronic transplantation.

### Transcriptomic analysis shows a composite of biological age modulation subsequent to age-disparate transplantation

To further understand the results of our DNAm analysis, we sequenced the transcriptomes of transplanted hearts. Differential expression analysis comparing young and old sham controls identified >100 significantly differentially expressed genes (DEGs), representing age-dependent changes in the mouse transcriptome. Surprisingly, few to no DEGs were identified in native hearts of heterochronic transplant recipients, whereas many DEGs were found in heterochronic vs. isochronic graft hearts; a similar pattern was observed in graft vs. native hearts from the same animals (**Figure 3A**).

**Figure 3.**
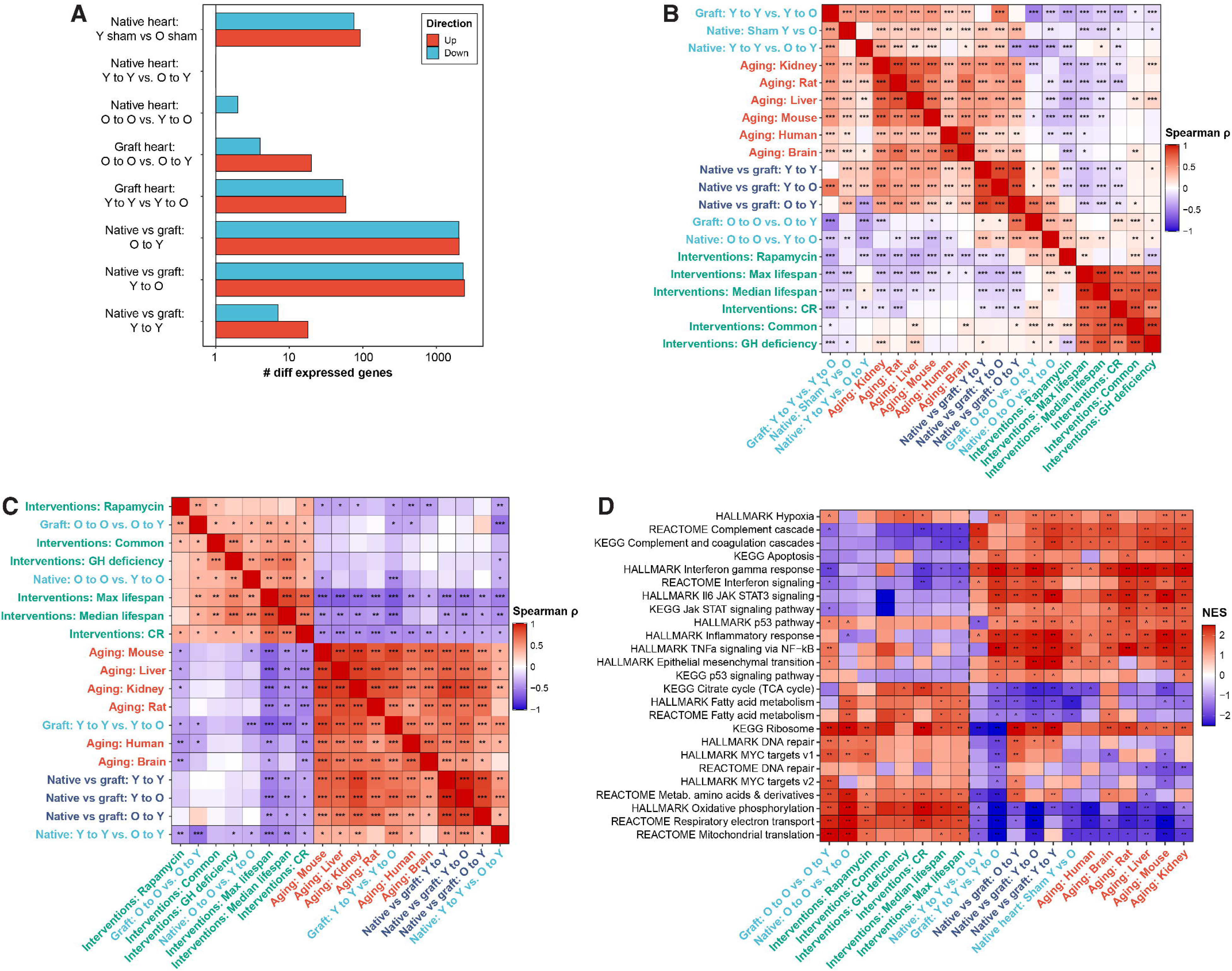
Transcriptomic signatures of aging support biological age assimilation in heterochronic heart grafts. (A) Number of significantly differentially expressed genes in various comparisons performed. (B–D) Associations between transcriptomic profiles of transplanted and native hearts and signatures of aging and longevity analyzed at the level of genes (B), cellular functions (C), and pathways (D). Spearman correlation and p value are shown. Sample sizes: Y sham native heart, Y to O native heart, and Y to O graft heart, n = 4; O to O graft heart, n = 6; n=5 for all other groups. Y, young; O, old.

We then applied transcriptomic signatures of aging and longevity^14,15^ to these data. These signatures were derived from meta-analyses of gene expression changes associated with aging and with lifespan-extending interventions across multiple tissues and species, yielding a defined set of genes consistently altered in the pro-aging direction and a complementary set consistently altered in the anti-aging, pro-longevity direction. When we examined the resulting data based on the analysis at both gene and pathway levels, we observed two clusters: (1) a pro-aging cluster, which positively correlated with signatures of aging and negatively correlated with signatures of anti-aging interventions, including: (i) old vs. young sham surgery animals, (ii) native hearts of young recipients with either heterochronic vs. isochronic heart transplants; (iii) young heterochronic vs. young isochronic heart transplants; and (iv) all transplants vs. native hearts from the same animals; (2) an anti-aging cluster, which negatively correlated with aging and positively correlated with signatures of anti-aging interventions, including: (i) old heterochronic vs. old isochronic heart transplants; and (ii) native hearts of young heterochronic hTx recipients (**Figure 3B–C**). These data were in general consistent with our DNAm clock analysis indicating that heart transplantation induced (i) pro-aging changes in young hearts grafted into older environments; (ii) anti-aging changes in old hearts grafted into young environments; and (iii), some degree of general pro-aging molecular changes in all graft hearts.

Thus, we conclude that the systemic environment strongly impacts the predicted biological age of heart transplants while the transplanted tissue exerts subtle to minimal effects on systemic biological age in our analysis.

### Functional enrichment analysis reveals factors associated with biological age modulation in heart transplants

We next performed functional enrichment analysis of our transcriptomic data to understand cellular pathways responsible for age modulation in our transplant model. Confirming our results thus far, we observed a clear separation of the data into the same clusters found previously. Notably, inflammatory pathways were upregulated in all heart transplants, consistent with a general pro-aging effect of the transplant procedure. Notably, several pathways were differentially modulated upon heterochronic transplantation. Interferon gamma response and interferon signaling pathways showed a mirrored, bidirectional association with these signatures, positively correlated in young heterochronic heart transplants and negatively in old heterochronic grafts. This pattern is consistent with the generic transplant effect noted previously rather than a signal specific effect to either direction of age mismatch. By contrast, the broader Hallmark inflammatory response and IL6/JAK/STAT3 signaling gene sets were significantly associated with aging signatures specifically in young heterochronic transplants but not in old heterochronically transplanted hearts. Mitochondrial-related pathways, including mitochondrial translation, oxidative phosphorylation, and respiratory electron transport, showed the most striking results, with large-magnitude, significant changes in both directions: downregulated in young heterochronic graft hearts and upregulated in old heterochronic graft hearts (**Figure 3D; Table S3**).

In summary, our experimental data demonstrate strong effects of heterochronic transplantation on the biological age of hearts, with the most prominent being an assimilation of transplant biological age driven by the age of the recipient.

### Human Cardiac Transplants assimilate recipient biological age at the molecular level

A key question that follows from our mouse data is whether similar biological age dynamics are at play in human recipients of heterochronic transplants. Our analysis of clinical heart transplantation revealed a median age disparity between donor and recipient (i.e., recipient age – donor age) of approximately +20 years (**Figure 4A**), driven by the increased need of older recipients **(Figure 4B**) and the preference for young donor organs (**Figure 4C**). We also found a smaller but non-negligible number of cases performed where the heart donor was considerably older than the recipient. To assess the biological age of the transplanted hearts in a subset of these patients, we gained access to historical endomyocardial biopsy samples from pathology archives, isolated DNA, and analyzed DNAm profiles using the HorvathMammalMethyl40 array.

**Figure 4.**
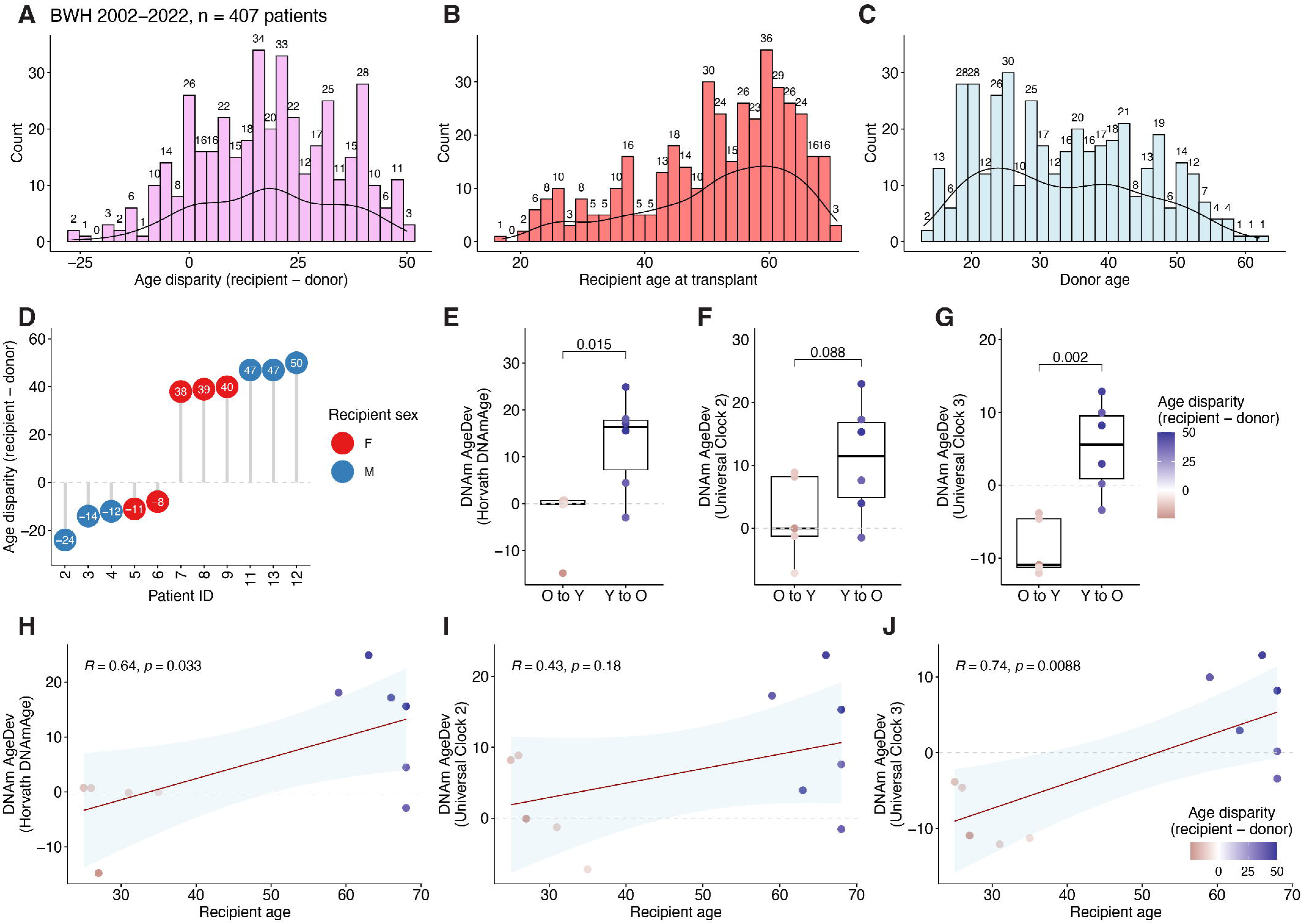
Heterochronic human heart transplants assimilate the recipient’s age. (A–C) Frequency of heterochronic heart transplants at Brigham and Women’s Hospital from 2002–2022. Counts for age disparity (A), recipient age (B) and donor age (C) are shown. (D) Demographic data of the heterochronic heart transplant cohort analyzed in our study. (E–G) DNAm AgeDev for endomyocardial biopsies from heterochronic heart transplant recipients calculated using the Horvath DNAm pan-tissue clock (E) and the universal mammalian clocks 2 (F) and 3 (G). p values were calculated with student’s unpaired t test. (H–J) Correlation between DNAm AgeDev and recipient chronological age. Pearson correlation and associated p value are shown. Sample sizes: O to Y, n = 5; Y to O, n = 6. Y, young; O, old.

We selected a group of 11 patients with age disparity between −24 and +50 years (**Figure 4D**) and examined AgeDev of the transplanted heart with respect to donor age.

Using the Horvath multi-tissue clock,^16^ we observed a clear and statistically significant difference in AgeDev between recipients of O (old) to Y (young) and Y to O heart transplants (**Figure 4E**). This difference was recapitulated using both Universal Clocks 2 and 3, although only the latter rose to the level of statistical significance (**Figure 4F–G**). Similarly, we observed significant positive correlations between AgeDev as determined by the Horvath multi-tissue clock and Universal Clock 3 and recipient age, indicating that the transplanted heart assumes the biological age of the recipient (**Figure 4H–J**). Thus, transplanted human hearts assimilate the biological age of their novel environment.

### Functional cardiac outcomes in human recipients confirm the clinical relevance of biological age assimilation

To determine whether biological age assimilation confers clinically relevant functional consequences, we analyzed detailed cardiac outcomes from electrocardiographic, echocardiographic, and cardiopulmonary stress test parameters collected at one-year post-transplant from hundreds of recipients at our center (**Table S4**). For each parameter, we modeled the independent association of recipient age with the outcome, adjusting for donor age and recipient sex. To allow direct comparison across parameters measured in different units and scales, we computed standardized coefficients, reflecting the change in each outcome, expressed in standard deviation units of that outcome, associated with a one standard deviation increase in recipient age (**Figure 5A-C** **and Table S5**).

**Figure 5.**
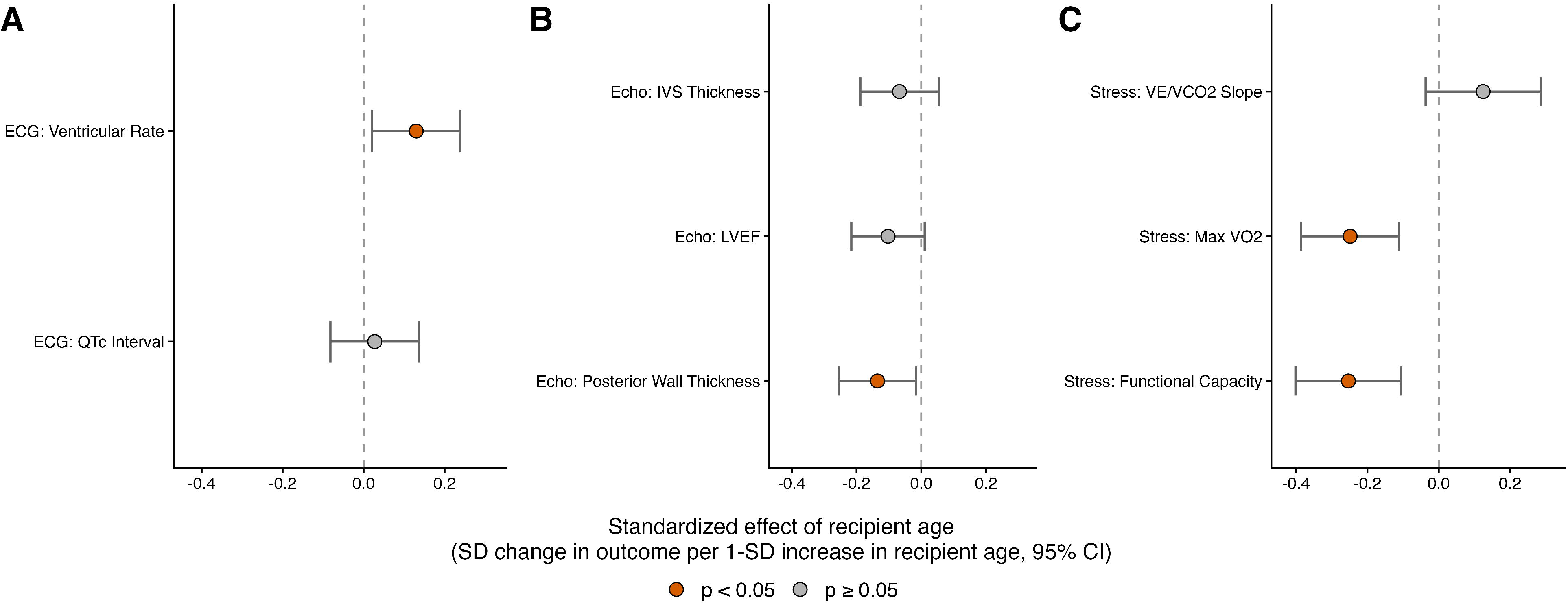
Functional cardiac outcomes in human heart transplant recipients are associated with recipient but not donor age. Standardized coefficients reflecting the independent association of recipient age with electrocardiographic (A), echocardiographic (B), and cardiopulmonary stress test (C) parameters, measured by one-year post-transplant in hundreds of recipients (cohort characteristics in **Table S4**). Coefficients reflect the change in each outcome, in standard deviation units of that outcome, associated with a one standard deviation increase in recipient age, adjusting for donor age and recipient sex. Sample sizes: (A) Ventricular rate (n = 317) and corrected QT interval (n = 317). (B) Left ventricular ejection fraction (n = 312), interventricular septal thickness (n = 272), and posterior wall thickness (n = 275). (C) Functional capacity (n = 162), maximal oxygen consumption (VO∼2∼ max, n = 151), and VE/VCO2 slope (ventilatory efficiency, n = 157).

Among electrocardiographic parameters, ventricular rate was significantly associated with recipient age, while the corrected QT interval was not (**Figure 5A**). Within echocardiographic parameters, posterior wall thickness was significantly associated with recipient age, while left ventricular ejection fraction showed a similar trend that did not reach significance, with interventricular septal thickness showing no association (**Figure 5B**). Most notably, recipient age was strongly and consistently associated with measures of exercise capacity: functional capacity and maximal oxygen consumption (VO_2_ max), both of which decline with normal chronological aging,^17^ were significantly lower with increasing recipient age, independent of donor age (functional capacity, p = 9.3×10-4, n = 162; VO_2_ max, p = 4.8×10-4, n = 151), while ventilatory efficiency showed no significant association (**Figure 5C**). The association between recipient age and these two exercise capacity measures remained significant after additional adjustment for BMI (**Table S5**).

Together, these data indicate that recipient age, rather than donor age, is associated with functional cardiac outcomes across multiple independent clinical domains, providing clinical validation of the molecular age assimilation described above at far greater scale than our endomyocardial biopsy cohort.

## Discussion

Our data demonstrate that systemic environment governs tissue biological age. These results broadly reinforce the idea that biological age may be affected by exposure to a novel systemic environment. In contrast to previous studies in heterochronic parabiosis models, we were able to probe the effects of transferring a single organ from its original environment to one of differing age. We found that the biological age of heterochronic heart grafts in mice is strongly affected by the age of the recipient while reciprocal effects on the recipient’s systemic biological age appeared minimal. This is in contrast to our previously published heterochronic parabiosis data,^7,8^ where reciprocal, systemic exchange of biological age was observed between parabionts (i.e. all examined old organs became biologically younger, and *vice versa*).

Strikingly, we found that similar biological age assimilation occurs in transplanted human hearts. In human patients, we found a strong association of the biological age of the transplanted hearts with the age of the recipients. This observation was particularly surprising given the large chronological age disparities between donors and recipients in the profiled cohort. Cardiac transplantation in humans occurs in a vastly more complex context compared to our standardized experimental mouse model, which utilized heterotopic transplants between healthy, inbred mice, minimizing the risks of allograft rejection and failure. Clinical heart transplantation is the only lifesaving treatment for end-stage cardiac failure, but commonly occurs against a backdrop of additional co-morbidities that frequently lead to the patient’s death while waiting for an organ.^18^ In addition, patients are treated after transplantation with an array of medications including immunosuppressants.^19^ It is thus remarkable that in spite of these complexities, biological age assimilation occurs in transplanted human hearts. This finding immediately raises the possibility that distribution of the limited pool of transplantable hearts may be optimized by allocating older organs to younger recipients. Further research will thus be needed to test whether reduction of biological age in transplanted organs leads to comparable (or improved) clinical outcomes for human heterochronic transplant recipients, and whether these effects overcome negative outcomes generally associated with older organs.^20,21^

Our detailed molecular findings were corroborated by an independent analysis of clinical outcomes at far greater scale. In hundreds of transplant recipients, recipient age, rather than donor age, was significantly associated with cardiopulmonary functional capacity at one-year post-transplant, providing an orthogonal, clinically grounded evidence for biological age assimilation. Notably, this functional signal was not uniform: recipient age associated strongly and consistently with measures of exercise capacity, but only selectively with structural and electrical parameters. This pattern echoes our genome-wide DNAm analysis, in which the effect of the heterochronic environment on graft tissue was detectable using targeted aging biomarkers but by unsupervised genome-wide analysis. Together, these observations suggest that biological age assimilation, at both the molecular and functional level, affects select measures rather than manifesting uniformly across all of them. It needs to be noted that our functional data are limited to one-year post-transplant and thus do not capture longer-term consequences such as chronic allograft vasculopathy or mortality, which are likely driven by structural and immunological components that are plausibly distinct from the dynamic molecular and functional state we describe (e.g., pre-existing vascular damage in older donor organs and cumulative alloimmune injury). Whether the biological age of a graft modulates these longer-term, structurally driven risks remains an important question for future studies.

Our study also raises important issues for ongoing and future research on biological age and strategies to target aging. At the most fundamental level, these data broadly support the concept that molecular damage accumulation represents the essence of aging.^22^ Exchange of accumulated damage between the recipient and the graft following transplantation would simultaneously explain why old hearts are rapidly rejuvenated when placed into a young environment, which can effectively dilute damage, and why young hearts placed into an old environment, and thus exposed to higher levels of damage, rapidly age. Our transcriptomic pathway analysis offers an initial candidate mechanism for this exchange, with mitochondrial and metabolic pathways emerging as the most consistently altered among aging-associated processes, suggesting a starting point for future mechanistic dissection. Complementary work has shown that transplantation of old organs has also been shown to promote senescence in recipient tissues through augmented release of SASP factors, including mitochondrial DNA,^10^ consistent with mitochondria as a central node in age-related graft-recipient crosstalk. Our recapitulation of the biological age assimilation effect in humans enabled by universal biomarkers of aging lends further credence to this notion and substantiates the hypothesis that modulation of biological age by the systemic environment is a deeply conserved phenomenon across species. Moving to more practical considerations, as attention in the field is increasingly focused on tissue/organ-specific aging and rejuvenation strategies,^23^ our data and other preliminary reports^24,25^ underscore the importance of the broader systemic context that should not be ignored. For instance, future research may address whether targeted rejuvenation of a single organ can persist over time if it occurs within the context of an aged organism.

We acknowledge that this study is limited in some respects. First, our mouse model of heart heterotopic heart transplantation is not perfectly analogous to the orthotopic procedure in humans. Although the sustained presence of the native heart in this system provides a useful experimental control, it is clearly not a perfect model, but orthotopic heart transplant in mice is considerably technically challenging. Our mouse model also does not induce an alloimmune response following transplant and thus differs from clinical heart transplantation. Nevertheless, our clinical studies demonstrated the impact of the environment on organ age in the presence of immunosuppression. Conversely, our study of heart biopsy samples from patients was limited by the need to assess historical fixed and paraffin-embedded samples, which are not ideal for DNAm profiling. However, our carefully controlled mouse study paired with our extensively quality-controlled human DNAm data present compelling evidence supporting age assimilation of tissues placed into heterochronic systemic environments. Future work will be needed to understand the temporal dynamics of biological age assimilation in heart transplantation, including the persistence of the apparent rejuvenation and whether it is ultimately reflected in long-term hard clinical outcomes. While it seems likely that our findings in heart transplantation will also apply to other organ transplants, future studies will need to confirm this assumption.

In summary, this study advances our understanding of fundamental features of biological age, namely that systemic biological age influences the biological age of individual organs. As these dynamics also occur in humans, our study suggests potential directions for future organ allocation protocols for transplant, which may contribute to reducing the critical shortage of transplantable hearts and potentially other organ transplants.

## Acknowledgements

We thank Dr. Robert Padera (Brigham and Women’s Hospital) for extremely helpful discussions and guidance. We are also grateful to the BWH Pathology Specimen Locator Core and Bobby Brooke of the Epigenetic Clock Development Foundation. Funded by grants from the National Institute on Aging (to VNG and SGT), the Hevolution Foundation (to VNG, SGT, and JRP), and the Pablo and Almuead Legorretta Kidney Health Research Fund (to SGT). JRP was supported by the BWH Organ Design and Engineering Training Program, NIBIB grant 5T32EB016652-07.

