## Supplementary Appendix for "Transplanted hearts assimilate the recipient’s biological age"

Vadim N. Gladyshev, PhD^1*, #^

^1^ Division of Genetics, Department of Medicine, Brigham and Women’s Hospital, Harvard Medical School, Boston, MA, USA

^2^ Division of Transplant Surgery and Transplant Surgery Research Laboratory, Department of Surgery, Brigham and Women’s Hospital, Harvard Medical School, Boston, MA, USA

^3^ Altos Labs, San Diego, CA, USA

^4^ Department of Surgery, Charité, Berlin, Germany

^5^ Division of Cardiovascular Medicine, Department of Medicine, Brigham and Women’s Hospital, Harvard Medical School, Boston, MA, USA

^†^Equal contribution as first authors

^#^Corresponding authors

*Correspondence:

**Methods**

Mouse experiments

All mouse experiments were approved by the Mass General Brigham IACUC. C57Bl/6 Mice were obtained from Charles River Laboratories or the NIA aged rodent colony and acclimated to our animal facility for at least 48 h. Mice were maintained in a barrier facility in sterilized, ventilated cages and fed standard laboratory chow (LabDiet 5053) and reverse osmosis drinking water ad libitum and maintained on a 12h:12h light:dark cycle. Mice were generally housed socially (2-3 mice/cage). Mice were humanely euthanized at the conclusion of each experiment by CO2 exposure followed by cervical dislocation.

Mouse heterotopic heart transplantation

Heterotopic heart transplantation was carried out with a modified cuff technique as previously described.^1^ Mice were anesthetized and donor hearts were anastomosed to the recipient’s common carotid artery and internal jugular vein. Ischemia and anastomosis times were kept consistent between animals. Graft function was monitored by palpation following the procedure. Animals with failed grafts and animals who died following surgery were not analyzed. Animals were maintained for 4 (old and young recipients) or 6 (middle-age recipients) months post-transplant, after which they were humanely euthanized, and tissues were collected for analysis.

Human samples

This study was approved by the Mass General Brigham IRB (protocol number 2022P002490). From the records of cardiac transplantation cases performed at Brigham and Women’s Hospital from 2002-2022, we searched for heterochronic transplant recipients, defined here as those recipients with at least a 5-year difference in chronological age with their donor. From this pool of subjects, we selected 5 patients who received an older heart and six recipients who received a younger heart. Pathology records were searched for these patients and stored paraffin blocks containing endomyocardial biopsy samples collected as part of routine post-transplant care were requested from the BWH Pathology Specimen Locator Core.

Isolation of nucleic acids

DNA was isolated from mouse tissue/blood samples using either the DNeasy Blood and Tissue Kit (Qiagen) or the Chemagic360 system with the 10 mg tissue or 400 ul blood kit (Revvity). RNA was isolated from mouse tissues using the Chemagic360 system with the 10 mg RNA tissue kit (Revvity). In all cases, the manufacturer’s protocols were followed. Generally, ~100 μl of blood or ~10 mg of solid tissue was used as starting material. DNA/RNA was eluted into nuclease-free water and DNA was concentrated using a speedvac when necessary. Concentration of DNA/RNA samples was determined using the Qubit or Quantit dsDNA BR or RNA HS assay kits (Invitrogen). Isolated DNA was stored at –20°C and isolated RNA was stored at –80°C.

DNA methylation profiling

Methylation data was generated through the Epigenetic Clock Development Foundation. For this study, mouse samples were profiled with the HorvathMammalMethyl320 array and human samples were profiled with the HorvathMammalMethyl40 array at Akesogen. Samples were randomized to avoid introduction of batch/chip effects. All sample preparation/processing was carried out according to the Illumina kit protocols.

Gene expression profiling

Total RNA isolated as described above was checked for quality using an Agilent 2100 Bioanalyzer. Samples that passed QC were paired-end sequenced on an Illumina NovaSeq 6000 S4 with 150 bp read length.

**Quantification And Statistical Analysis**

Principal component analysis of genome-wide DNA methylation

Principal component analysis (PCA) was performed on genome-wide DNAm beta values from the mouse heart transplantation cohort, restricted to probes with non-missing beta values in every sample. PCA was computed using the R prcomp function with mean-centering and without variance scaling. Native hearts and graft hearts were analyzed as separate PCAs, each computed independently on the relevant sample subset.

DNAm clock analysis

For mammalian microarray analysis, raw methylation data were first normalized using the SeSAMe R package and beta values were calculated. Samples with anomalously low fractions of detected probes (<20%) were considered to have failed and were not analyzed. Samples that were determined to be significant outliers using principal component analysis were also excluded. DNAm age biomarkers were calculated as previously described.^2–4^ AgeDev was calculated by subtracting chronological age from the prediction. The DNAm data in this manuscript was generated in two batches. Because raw AgeDev values showed systematic inter-batch offsets, a within-batch ratio normalization relative to sham controls was applied prior to group comparisons. For each tissue-batch stratum, the minimum AgeDev value was identified and its absolute value added to all values in that stratum to ensure non-negativity; each shifted value was then divided by the mean shifted AgeDev of sham animals within the same stratum. This procedure sets the sham mean to 1.0 within each batch while preserving relative differences among experimental groups profiled together, removing batch-level technical variation without altering within-batch biological comparisons.

Differential methylation analysis

Differential methylation modeling was carried out using SeSAMe.^5^ Probes not significantly detected were excluded, as were SNP probes. Modeling was performed using the DML function. CpGs were considered significantly differentially methylated if Benjamini-Hochberg adjusted p value was less than 0.05 and the change in methylation level was at least 5%. These thresholds were chosen to be deliberately stringent, prioritizing robustness and reproducibility of the identified sites. Gene names for identified CpG sites were imported from the Illumina Infinium array manifest and used to label points in figures for clarity.

Genome-wide correlation of transplant-induced and intrinsic aging methylation changes

To test whether methylation changes induced by heterochronic transplantation were directionally concordant with intrinsic aging across the genome, without restricting to a significance-filtered CpG subset, per-CpG correlations were computed across all probes with non-missing beta values in every profiled sample. No probes were excluded on the basis of statistical significance or effect size in either axis. For each CpG, two quantities were calculated: the heterochronic delta-beta, defined as the mean beta difference between heterochronic and isochronic grafts (young-to-old graft minus young-to-young graft for the accelerated-aging direction; old-to-young graft minus old-to-old graft for the rejuvenation direction), and the intrinsic aging delta-beta, defined as the mean beta difference between native hearts of old and young sham animals. Pearson and Spearman correlations between the heterochronic delta-beta and the intrinsic aging delta-beta were computed genome-wide for each direction.

Gene expression analysis

For RNAseq data, we filtered out genes with low number of reads, keeping only the genes with at least 10 reads in at least 50% of the samples. Filtered data was then passed to RLE normalization.^6^ Differential expression of genes was analyzed using edgeR,^7^ comparing heterochronic graft hearts to isochronic graft hearts, heterochronic native hearts to isochronic native hearts, graft hearts to native hearts from the same animals, and old versus young sham-operated animals. Obtained p-values were adjusted for multiple comparison with Benjamini-Hochberg method.^8^

Association with gene expression signatures

Association of gene expression log-fold changes induced by heterochronic heart transplantation with previously established transcriptomic signatures of aging was examined as described previously.^9,10^ Multi-tissue mouse signatures obtained via a meta-analysis of age-related gene expression changes from multiple datasets were utilized for this analysis.

First, for every signature we specified 250 genes with the lowest p-values and divided them into up- and downregulated genes. These lists were subsequently considered as gene sets. Then, we ranked genes differentially expressed in each comparison of interest (heterochronic versus isochronic graft hearts, heterochronic versus isochronic native hearts, graft versus native hearts, and old versus young sham-operated animals) based on their p-values, calculated as described above. Afterwards, we utilized gene set enrichment analysis (GSEA)^11^ to calculate normalized enrichment scores (NES) separately for up- and downregulated lists of gene sets as described in,^9^ and calculated the final NES as a mean of the two. To calculate statistical significance of obtained NES, we performed permutation testing where we randomly assigned genes to the lists of gene sets, maintaining their size. To get the p-value of the association between each transplant comparison and a certain signature, we used 5,000 permutations and calculated the frequency of random final NES that are larger in magnitude than the observed final NES. To adjust for multiple testing, we performed a Benjamini-Hochberg correction. Final NES for association of each transplant comparison with aging signatures were used to generate figures.

Functional enrichment analysis

For the identification of enriched functions distinguishing isochronic and heterochronic mice, we performed functional GSEA^11^ on a pre-ranked list of genes based on log_10_(p-value) corrected by the sign of regulation, calculated as:

$-\left( pv \right)\times sgn\left( lfc \right)$,

where *pv* and *lfc* are p-value and logFC of a certain gene, respectively, obtained from edgeR output, and *sgn* is the signum function (equal to 1, -1 and 0 if value is positive, negative or equal to 0, respectively). REACTOME, KEGG and HALLMARK ontologies from the Molecular Signature Database (MSigDB) were used as gene sets for GSEA. The GSEA algorithm was performed separately for each comparison of interest (heterochronic versus isochronic graft hearts, heterochronic versus isochronic native hearts, graft versus native hearts, and old versus young sham-operated animals) via the *fgsea* package in R with 5000 permutations. A q-value cutoff of 0.1 was used to select statistically significant functions.

Similar analysis was performed for gene expression signatures of aging. Pairwise Spearman correlation was calculated between the NES of each transplant comparison and the NES of each aging/longevity signature. A heatmap colored by NES was built for manually chosen statistically significant functions (adjusted p-value < 0.1). Complete list of functions enriched at least one transplant comparison is included in Table S3.

Clinical outcomes analysis

To test whether recipient biological age, rather than donor heart age, predicts post-transplant cardiac function, cross-sectional linear regression analyses were performed at one-year post-transplantation using electrocardiographic, echocardiographic, and cardiopulmonary exercise testing (stress test) data collected from electronic medical records. For each continuous outcome, a linear regression model was fit with recipient age, donor age, and recipient sex as covariates. For stress test-derived outcomes, an additional sensitivity model further adjusted for BMI, available for a subset of stress test participants. Coefficients for recipient age are reported as standardized (SD-unit) effect sizes with 95% confidence intervals. Sample sizes varied by outcome due to differing data availability across parameters and are reported in the corresponding figure legend.

Statistics

In general, for group comparisons, ANOVA was first used to test for significant variance between groups. If these tests revealed a significant effect, paired t tests corrected by controlling the false discovery rate using the Benjamini-Hochberg method were carried out between groups. Exact p values are shown within all figures. For single comparisons, unpaired student’s t test was used. Correlation was tested with either Pearson or Spearman correlation as indicated in figures. All t tests were two-tailed. Sample sizes are indicated in figure legends.

### **Supplementary Figure**

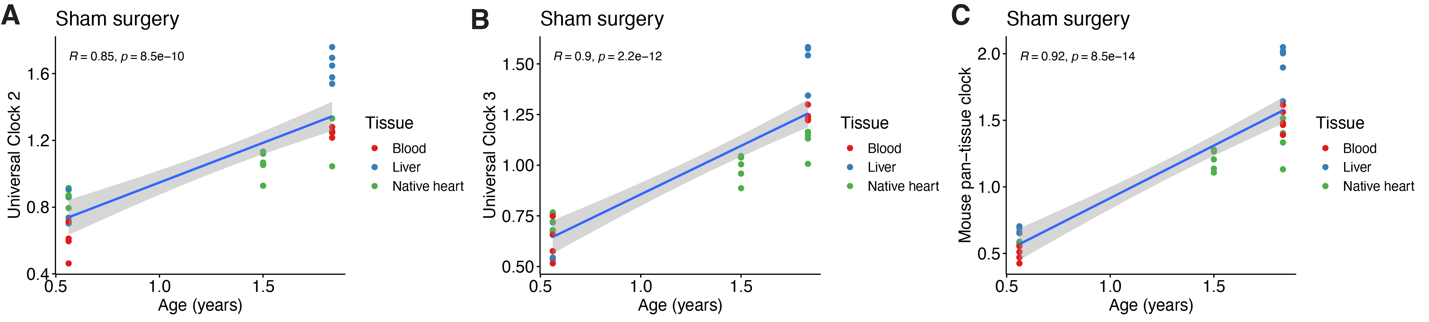

**Figure S1, related to Figure 1. Correlations between age predictions and chronological age for animals subjected to sham surgery.** DNAm age predictions calculated with Universal Clock 2 (A), Universal Clock 3 (B), and the mouse pan-tissue clock (C). Pearson correlation coefficient is shown.

### **Supplementary Tables**

**Table S1, related to Figure 1. Full statistical comparison results.** See .csv file.

**Table S2, related to Figure 2. Results of differential methylation analysis.** See .xlsx file.

**Table S3, related to Figure 3. Results of gene set enrichment analysis of transcriptomic data.** See .xlsx file.

**Table S4, related to Figure 5. Functional cohort baseline characteristics.** See .csv file. N reflects the total number of patients with a recorded measurement for that modality at one-year post-transplant. Analytic sample sizes vary by outcome within each modality owing to missing values in one or more model covariates (recipient age, donor age, recipient sex, BMI); outcome-specific Ns are reported in Table S5. Values reported as mean ± standard deviation, with values in parentheses representing the observed range (minimum–maximum).

**Table S5, related to Figure 5. Complete regression results for clinical functional outcomes, including raw and standardized coefficients and BMI-adjusted sensitivity analysis.** See .csv file.
